# Predicting the subjective intensity of imagined sensory experiences from electrophysiological measures of oscillatory brain activity

**DOI:** 10.1101/2023.10.31.564917

**Authors:** Derek H. Arnold, Blake W. Saurels, Natasha Anderson, Isabella Andresen, Dietrich S. Schwarzkopf

**Author notes:** **Data and materials availability** All EEG data and analysis scripts for this project will be made available via UQeSpace https://espace.library.uq.edu.au/. **Author contributions** D.H.A conceived of the study, programmed experiments, analysed data and wrote the first draft of the manuscript. I.A., N.A. & B.W.S. all tested participants and edited successive versions of the manuscript. D.S. provided advice on data analyses and edited successive versions of the manuscript.

## Abstract

Most people can conjure images and sounds that they experience in their minds. There are, however, marked individual differences. Some people report that they cannot generate imagined sensory experiences at all (aphantasics) and others report that they have unusually intense imagined experiences (hyper-phantasics). These individual differences have been linked to activity in sensory brain regions, driven by feedback. We would therefore expect imagined experiences to be associated with specific frequencies of oscillatory brain activity, as these can be a hallmark of neural interactions within and across regions of the brain. Replicating a number of other studies, relative to meditation we find that the act of engaging in imagining auditory or visual sensations is linked to reductions in the power of oscillatory brain activity across a broad range of frequencies, with prominent peaks in the alpha band (8-12 Hz). This oscillatory activity, however, did not predict individual differences in the subjective intensity of imagined experiences. For imagined audio experiences, these were rather predicted by reductions within the theta (6 – 9 Hz) and gamma (33 – 38 Hz) bands, and by increases in beta (15 – 17Hz) band activity. For imagined visual experiences these were predicted by reductions in lower (14 – 16Hz) and upper (29 – 32 Hz) beta band activity, and by an increase in mid-beta band (24 – 26 Hz) activity. Our data suggest that there is sufficient ground truth to the subjective reports that people use to describe the intensity of their imagined sensory experiences such that these can be predicted by the power of distinct rhythms of brain activity.

## Introduction

Most people can conjure images and sounds that they experience within their minds [1] – such that we can refer to the mind’s eye and to the mind’s ear. This capacity is so general that you might be surprised that some people have no such capacity. People who are unable to voluntarily form mental images are known as aphantasics [2]. These people often report also being unable to voluntarily generate and experience a mental soundscape, other than internal speech [3-4]. To an aphantasic, like the first author, the capacity of others to voluntarily generate images and sounds that they then experience within their minds can seem mysterious, leading them to ponder if most people have frequent hallucinations during their waking lives. We can each struggle to understand that the experiences of others might be unlike our own.

Aphantasics might be positioned at one end of a continuum, with people who are prone to have intense imagined sensory experiences at the other extremity [5]. Schizophrenics [6-7] and people with PTSD [8], for instance, tend to report having vivid imagined visual experiences.

We no longer need to rely entirely on self-report measures for evidence of individual differences in the intensity of imagined sensory experiences. The existence of subjective differences seems to have been substantiated by studies that have shown that the subjective intensity of imagined sensory experiences can modulate performance in an *objective* test of visual sensitivity [9-11]. Moreover, differences in the degree to which imagery modulates performance in an objective measure of visual processing has been linked to morphological differences between different human brains. Specifically, the degree of modulation has been found to be inversely scaled with the surface sizes of V1 (primary visual cortex) and V2 [12]. So, people with smaller V1s tend to report having more intense imagined experiences, and this predicts the degree to which imagery will modulate performance in an objective test of visual sensitivity [9].

Evidence linking imagined experiences to activity in sensory brain regions [e.g. 12] is consistent with the hypothesis that *endogenously* driven imagined sensory experiences are driven by a reversed processing hierarchy, relative to *exogenously* driven experiences [1,13]. Specifically, the theory is that imagined experiences are driven by a cascade of neural events, with processes originating in frontal brain regions triggering activity in medial and temporal brain regions, which enacts the retrieval of memories, which in turn triggers activity in primary sensory brain regions. Hypothetically, it is the activity in primary sensory brain regions, ultimately triggered by processes in frontal brain regions, that is causally involved in the generation of imagined sensory experiences [13-14]. According to this view, we would additionally expect imagined experiences to be associated with the power of specific frequencies of oscillatory brain activity, as these can serve as a hallmark of different cognitive operations and the interactions that they set in train between populations of neurons both within [15] and across different regions of the brain [15].

There are well established links between oscillatory brain activity and the generation of imagined sensory experiences. For instance, internally directed cognitive operations tend to be associated with reductions in the power of alpha band (8 – 12Hz) oscillatory brain activity [16-18], and there is good evidence that alpha-band power reductions are associated with the generation of imagined visual experiences [e.g. 19-21], but for contrary evidence see [22]. Indeed, recent evidence suggests that alpha-band oscillations can serve as a signature of shared visual mental imagery and perceptual representations [21]. This evidence, however, leaves open the question of whether alpha-band oscillatory brain activity in particular, or oscillatory brain activity in general, might also predict the subjective intensity of different peoples’ imagined sensory experiences.

Here, we test the hypothesis that the power of oscillatory brain activity can predict individual differences in the subjective intensity of imagined audio and visual experiences. To preface our results, we first demonstrate compliance with task instructions (that participants should conjure different types of imagined sensory experience – audio or visual, or that they should meditate). This is achieved by showing that each of these operations is characterised by distinct patterns of oscillatory brain activity, that are both reliable across participants, and are consistent with patterns of brain activity identified in previous studies. Moreover, we can decode what type of cognitive operation a given participant was engaged in on a trial-by-trial basis, and we establish that this capacity does not correlate with the subjective intensities of different people’s imagined experiences. This is important, as it implies that people who described their imagined experiences as relatively weak or strong were equally engaged in our tasks, as we could equally discern what cognitive operation they had engaged in from analyses of their brain activity. Finally, we find that we can predict the subjective intensity of a person’s imagined visual experiences from analyses of the oscillatory power of their brain activity. In our sample, intense imagined audio experiences were predicted by reductions in the theta (6 – 9 Hz) and gamma (33 – 38 Hz) bands, and by increases in beta (15 – 17 Hz) band activity. For visual imagery intense imagined visual experiences were predicted by reductions in lower (14 – 16 Hz) and upper (29 – 32 Hz) beta band activity, and by an increase in mid-beta band (24 – 26 Hz) activity.

## Methods

### Ethics

Ethical approval was obtained from the University of Queensland’s (UQ) Ethics Committee, and the experiment was performed in accordance with UQ guidelines and regulations for research involving human participants. Each participant provided informed consent to participate in the study and were made aware that they could withdraw from the study at any moment without prejudice or penalty.

### Participants

Forty nine people volunteered to participate in the study. These included one of the authors (IA), and 48 undergraduate students and staff at The University of Queensland (40 female, M_age_ = 22.2). This should be regarded as a convenience sample. Testing took place during a period marked by intermittent disruptions of face-to-face testing, so we were unable to obtain our desired sample size (of 100), which would have delivered ∼0.97 power to detect medium sized (∼0.39) linear relationships between behavioural and neural measures. Our obtained sample size delivers an equivalent power to detect strong sized (∼0.59) effects. While this research was carefully planned, study procedures and analyses were not pre-registered.

Equipment failure (event triggers failed to register) resulted in data for 5 participants being unusable. The final group of participants, who contributed data that was analysed, therefore consisted of 44 people, including one author and 43 undergraduate students and staff (35 female, M_age_ = 22.2). We included an author in data collection as our hypotheses relate to the power of oscillatory brain activity, distributed across the brain and frequency spectrum, and we felt it was improbable that foreknowledge of this interest could favour a particular outcome – especially as no neuro-feedback was provided during recordings of brain activity.

### Apparatus and stimuli

Testing took place in a darkened room. A chinrest was used to ensure a constant distance (50 cm) from the monitor, and to stabilize the head during EEG recordings. Trial instructions were presented on an ASUS VG248QE 3D Monitor (1920 x 1080 pixels, refresh rate: 60 Hz), driven by a Cambridge Research Systems ViSaGe stimulus generator and custom MATLAB R2015b software. A Tucker-Davis Technologies (TDT) Audio Workstation was used to produce auditory white noise, which was emitted diotically at a clearly supra-threshold intensity (∼50 dB SPL) by speakers positioned to either side of the testing display. EEG data were recorded using a Biosemi International ActiveTwo system. Electrodes (64 Ag/AgCl) were placed according to the extended international 10-20 system and digitised at a 1024 Hz sample rate with 24-bit analog-digital conversion. The standard BioSemi reference and ground electrodes were used during recordings.

All data and analysis scripts underlying the results on which study conclusions are based are available to via UQ eSpace (analysis scripts: https://doi.org/10.48610/b09a867 Imagination spectra files: https://doi.org/10.48610/a1b3395 Meditation Spectra files: https://doi.org/10.48610/59e5757).

### Procedure

Trial sequences are depicted in Figure 1. Before all trials, there were written instructions on the test display stating “On this trial, you should close your eyes before you left click to begin the trial. This will trigger an audio white noise presentation. Please keep your eyes closed until the noise stops”. On meditation trials, participants were additionally instructed to “try to empty your mind, relax, and concentrate on your breathing. Try to ignore any visual or auditory experiences until the noise stops”. On Audio Imagination trials, participants were then asked “While you have your eyes closed, please try to create a mental soundtrack”. Five instructed scenarios were selected at random, including asking people to imagine hearing a favourite piece of music, the sounds of a busy street, the sound of their childhood caregiver, the sound of birds singing, or the sounds of a summer thunderstorm (which are common in Brisbane, Australia – the site of research). On Visual trials, people were asked to try and create a mental image, of either their childhood caregivers face, of themselves waiting to cross a road, a sunrise over water, a duck landing on a lake, or of their loungeroom. Before all trials, instructions finished with the directive “When you are ready to start the trial, close your eyes and click the left mouse button”.

**Figure 1.**
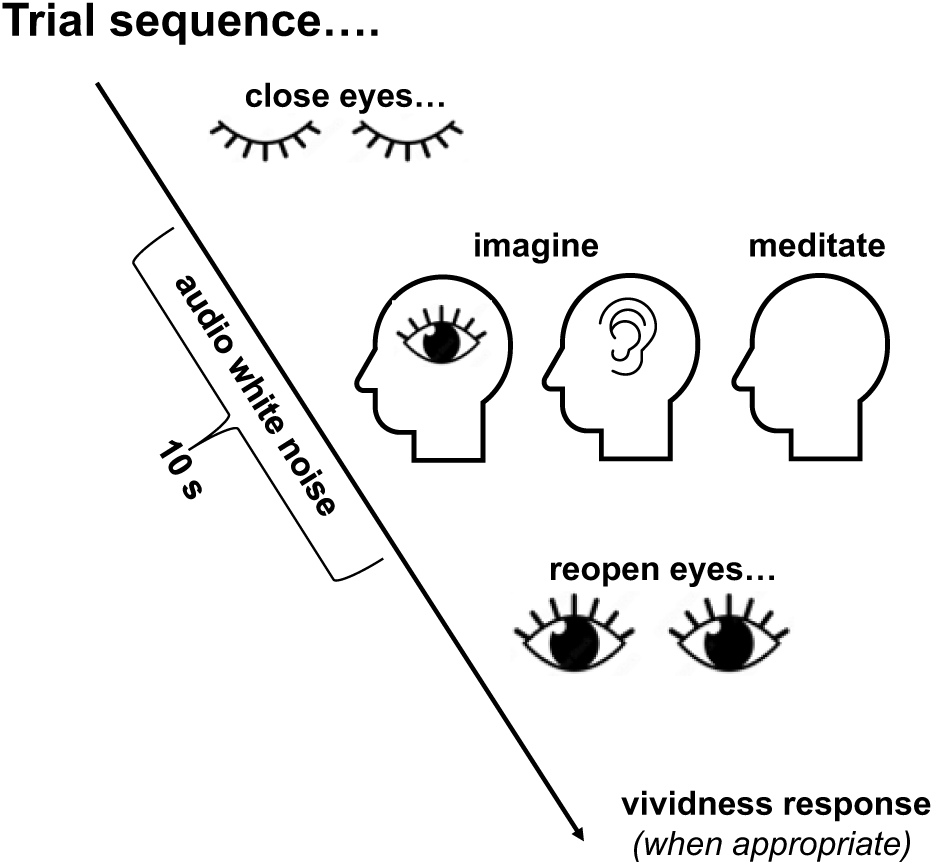
Graphic depicting a trial sequence. Each trial began with participants reading trial task instructions. They would then close their eyes before pressing a mouse button to begin the trial. After a short delay (0.62-1.62 seconds), this initiated a 10 second period wherein soft white noise was presented via speakers. On Audio and Visual Imagination trials, during these periods participants tried to imagine having a sensory experience in the specified sensory modality. On meditation trials, participants tried to empty their minds and concentrate on their breathing. In each case, participants were prompted to re-open their eyes by the white audio noise stopping. On Audio and Visual Imagination trials, they would then rate the vividness of their imagined experience.

After Imagination Condition trials, participants were asked to indicate how vivid their imagined sensory experiences had been, using a 5-point scale (from “No image / mental soundtrack at all. I only know that I was thinking of images / sounds”, to “Perfectly realistic, as vivid as real seeing / as if I was listening to my mental soundscape”). These responses were adopted from the vividness of visual imagery questionnaire (VVIQ2) [23]. A full experimental session involved 54 individual trials, 18 for each of the 3 experimental conditions, all interleaved in a random order. This number of trials may seem small, but the reader should be mindful that the individual trials were protracted (each taking ∼20 seconds to complete, and encompassing 10 seconds of sustained sensory imagination, or meditation), and they therefore generated a considerable amount of brain imagining data per individual trial.

### EEG data pre-processing and analyses

All analyses of data were conducted using custom MATLAB scripts using the FieldTrip toolbox [24], and MATLAB’s in-built commands Fast-Fourier transform (fft) and Fit Regression Support Vector Machine (fitsvrm). FieldTrip routines were used to high- (1 Hz), low- (100 Hz), and notch (45-55 Hz) filtered EEG data using a 6^th^ order Butterworth filter with a two-pass direction. Data for each sensor were then re-referenced to the volume average voltage (from across all 64 channels). Eye and muscle artifacts were detected via an independent-components analysis (ICA) and removed. Data was then epoched into 9 second segments, centred on a 10-second trial sequence (see Figure 1). This epoch selection avoids periods contaminated by transients relating to people closing or reopening the eyes. Data were then sorted into Audio Imagination, Visual Imagination and into Meditation epochs. For all epochs, spectra were calculated for data from all 64 channels using the Matlab fft command. The range of amplitudes recorded by each sensor during an epoch was first checked, and if this exceeded 250mV trial data for that sensor was excluded from analyses (by setting the power estimate to NaN). Otherwise, the matlab fft command was used to calculate an estimate of oscillatory power for each frequency. These were averaged across 1Hz bins, providing rounded estimates of oscillatory power from 1 to 40Hz, for each sensor on each trial for each participant.

## Results

### Correlations between the intensities of imagined Audio and Visual experiences

We examined relationships between average individual ratings of the intensity of imagined audio and visual experiences. A Pearson’s correlation revealed a robust positive correlation (r = 0.70, p < 0.001, see Figure 2). These data are consistent with datasets demonstrating similarly strong positive correlations between the subjective intensity of imagined audio and visual experiences [e.g. 4].

**Figure 2.**
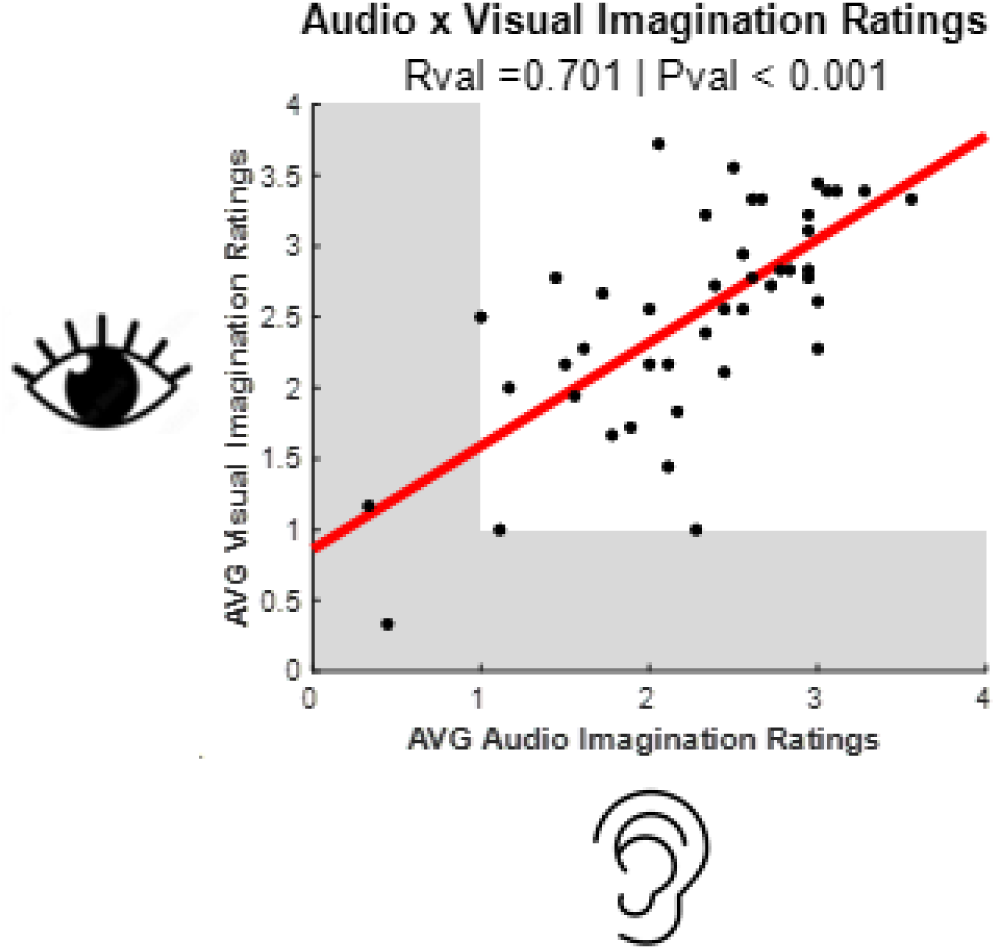
Scatter plot of average Visual (Y-axis) and Auditory (X-axis) imagination intensities reported by each participant. Shaded grey mark regions where scores might be taken as evidence for aphantasia (i.e. these scores indicate that participants *always* rated imagined sensory experiences as “No image / mental soundtrack at all. I only know that I was thinking of images / sounds” or as “Vague and not at all clear”.

### Conditional Spectra

As all recordings of brain activity were taken while people had their eyes closed, across all conditions there were prominent peaks in estimates of the power of alpha-band (∼10Hz) oscillatory brain activity (see Figure 3). This was true whether oscillatory power was averaged across all sensors (see Figure 3, top row) or was only estimated from a subset of occipital / parietal sensors (sensors PO7, PO3, O1, PO8, PO4 and 02; see Figure 3, bottom row).

**Figure 3.**
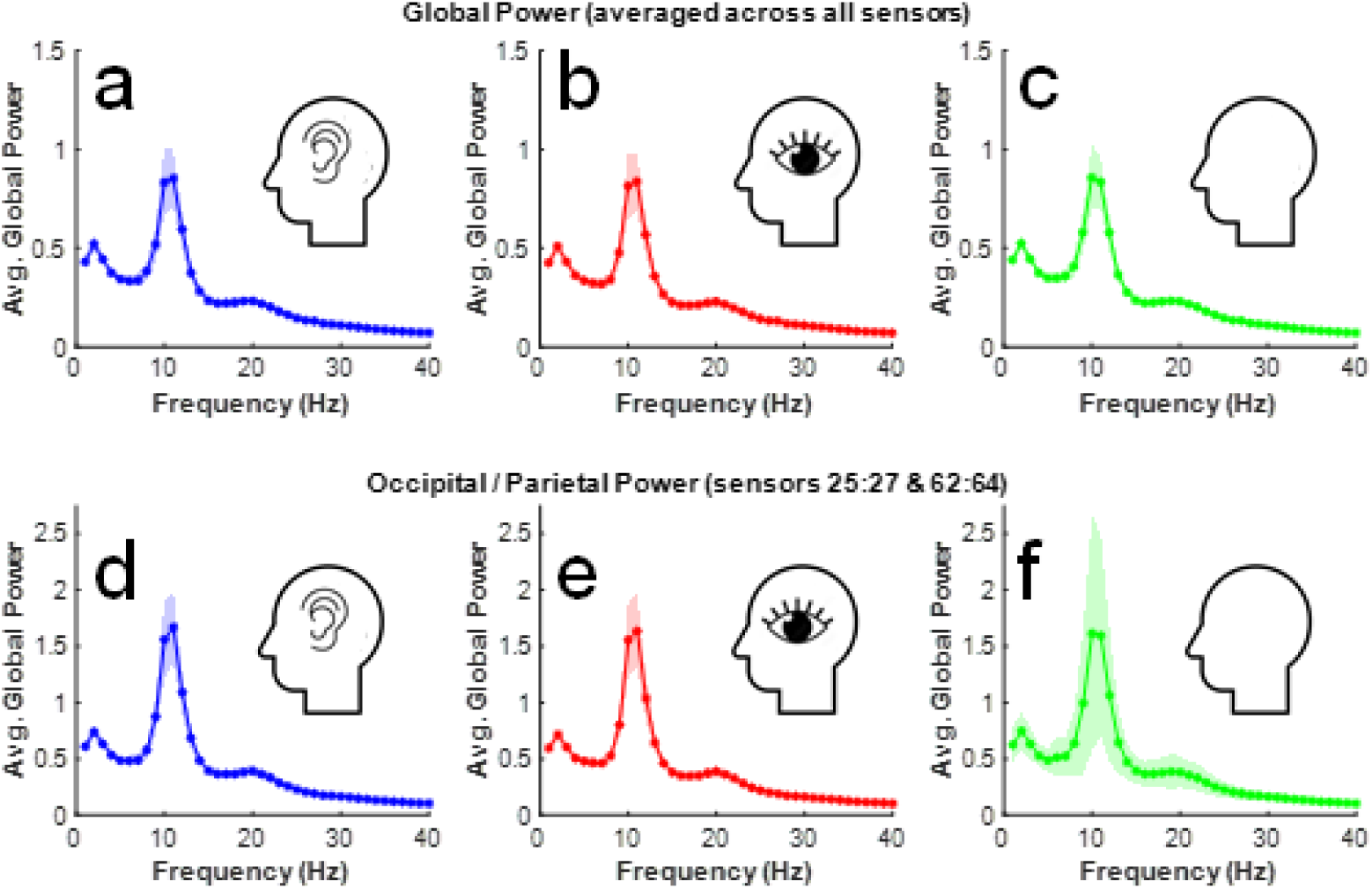
Plots of estimates of oscillatory power (Y-axes) as a function of frequency (X-axes). These data are averaged across all participants, separately for each condition. In top panels (**a-c**) data are averaged across all sensors. In bottom panels (**a-c**) data are averaged across occipital / parietal sensors (sensors 24:26 and 62:64). Data recorded while people imagined having audio experiences are depicted in panels **a** and **d**. Data recorded while people imagined having visual experiences are depicted in panels **b** and **e**. Data recorded while people meditated are depicted in panels **c** and **f**.

### Trial-by-trial decoding of experimental condition for individual participants

A reasonable concern would be that our participants might not have followed task instructions, to imagine having audio or visual experiences, or to meditate on designated trials. To confirm that our participants were likely following task instructions, we conducted a trial-by-trial decoding process for each participant, based on a nearest neighbour classification process with jack-knifed cross validation [e.g. 16, 25]. In these analyses trial data were only ever compared to other trials from the same participant.

For these analyses, estimates of oscillatory power (from 1 to 40 Hz) for each sensor (1 – 64) were used as features in a decoding process, so the feature space for each participant was a 64 sensor X 40 Hz spectrum of oscillatory power estimates X 3 conditions (Imagine Audio, Imagine Visual, and Meditation). The individual classification process was a leave-one-trial-out cross validation scheme, which is essentially a test for reliably distinct spectra prevailing on trials for each of our experimental conditions. For each trial training sets consisted of data from all other trials. Conditional signatures were calculated by averaging training set data across all trials from the same condition, and the left out trial data was decoded as having belonged to the condition with which the sum of absolute conditional residuals (a sum of unsigned difference scores) was smallest. Note that circularity was avoided by excluding the to-be-decoded trial data from the training set. Decoding was successful when the to-be-decoded trial data was matched to the experimental condition to which it belonged – due to having the smallest sum of unsigned residuals with that conditional signature. Note that simple absolute difference scores were used for this determination, as opposed to calculating a Euclidean distance, and no rescaling of data was conducted. Decoding success rates are calculated as the proportion of correctly classified trials. Chance decoding success rates are ∼0.333, as there were three conditions.

In Figure 4a we have plotted individual decoding success rates as a function of the average ratings used by participants to describe the subjective intensities of their imagined visual experiences. Note that overall, decoding success rates were well above chance (the bold horizontal black line marks the chance decoding success rate; single sample t_42_ = 10.13, p < 0.001), with a Bayes Factor analysis (BF10 > 10,000) revealing extreme evidence for the alternative hypothesis, that individual trial-by-trial decoding success rates would be above chance. All Bayes Factor analyses were conducted using the BayesFactor toolbox for Matlab [26].

**Figure 4.**
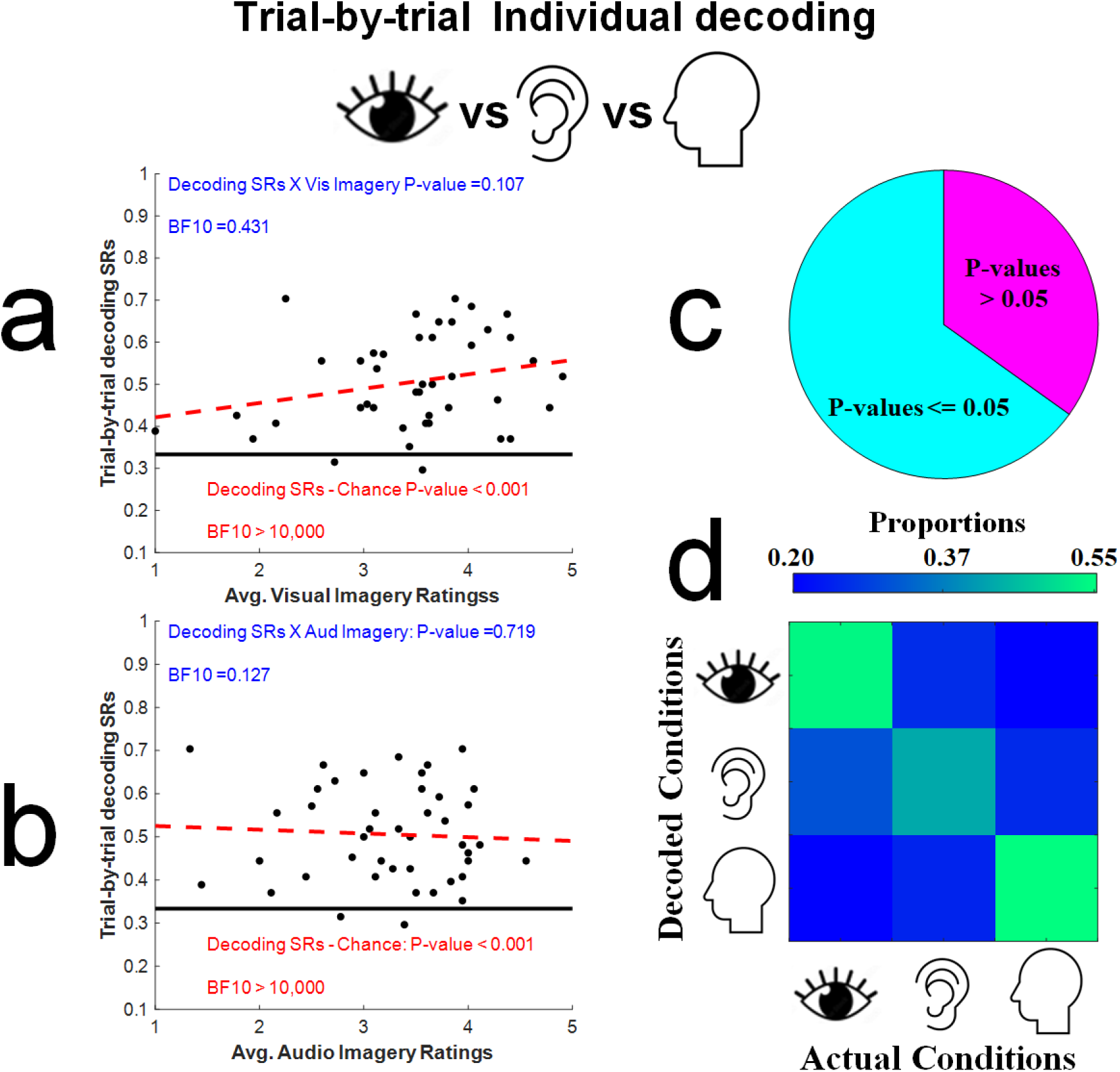
**a)** Scatterplot of trial-by-trial decoding success rates, calculated for each participant (Y-axis), and the average subjective ratings given to imagined experiences on visual imagination trials (X-axis). Blue text relates to statistical tests for a correlation between trial-by-trial decoding success rates and the subjective intensity of imagined experiences, whereas red text relates to statistical tests comparing actual decoding success rates to a chance decoding success rate (33.3%, marked by the black horizontal line). **b)** Details are as for Figure 4a, but analyses relate to average subjective ratings given to imagined experiences on audio imagination trials. **c)** Pie chart showing the proportion of participants for whom the experimental condition (imagined visual, audio or meditation trials) could be decoded on a trial-by-trial basis, from analyses of the spectra of their brain activity, at a rate that was statistically above chance (established via a non-parametric shuffle test, see main text for further details). **d)** Decoding Confusion Matrix, depicting proportions of decoded trials averaged across participants as a colour map, X-axis denotes actual experimental conditions and Y-axis the decoded conditions.

There was no robust relationship between decoding success rates and the subjective intensities of imagined visual experiences (Pearson’s r_42_ = 0.25, p = 0.11). We take this as evidence that people who had reported having relatively strong or weak imagined visual experiences were equally likely to have been following task instructions, as overall we could decode the experimental condition on a trial-by-trial basis with equal success from all participants, regardless of the ratings they used to describe the subjective intensities of their imagined visual experiences. Similar data are plotted in Figure 4b, for analyses relating to the subjective intensities of imagined audio experiences (Pearson’s r_42_ = -0.06, p = 0.72).

Our decoding process delivered a P-value relating to each participants’ decoding success rate, calculated via a non-parametric shuffle test. A P-value < 0.05 indicates that when decoded trial experimental conditions were randomly reassigned to experimental conditions 1000 times, less than 5% of the shuffled decodings resulted in a greater or in a matched success rate relative to our actual decoding process. As depicted in Figure 4c, we achieved decoding P-values of < 0.05 for 67% of our participants.

### Cluster tests for consistent conditional spectra differences across participants

We also conducted non-parametric cluster-based permutation tests, based on differences in the spectra of brain activity across different experimental conditions (see Figures 5-6). For these tests oscillatory power estimates were first averaged across trials for each participant, separately for each experimental condition. So, the feature space informing these tests was a 44 participant X 64 sensor X 40 Hz spectrum of oscillatory power estimates X 3 experimental conditions. For each sensor X frequency X condition combination, outlier features were identified (i.e. features > +/- 3 S.D.s from the conditional mean). When an outlier was detected, the individual’s feature estimate was replaced by a value interpolated across oscillatory power estimates from neighbouring sensors for that participant, when that was possible. Otherwise, the outlier was excluded from analyses by settings its value to NaN.

**Figure 5.**
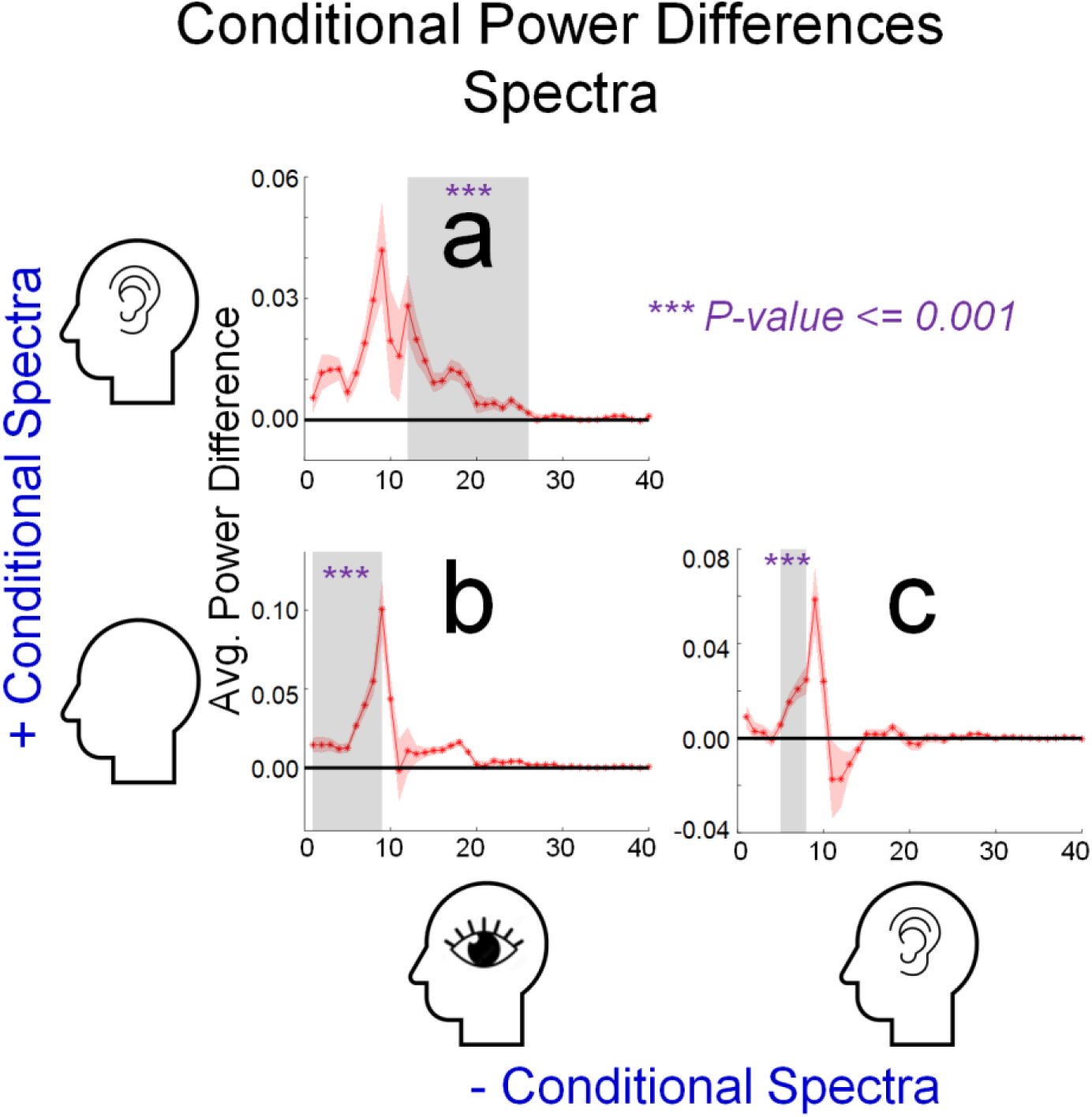
Plots depicting the results of significant non-parametric cluster tests, for differences in the power of oscillatory brain activity in different experimental conditions (Y-axes) as a function of frequency (Hz, X-axes). Red shaded regions of each plot depict +/- 1 SEM from the average difference between conditions. The frequency limits (Hz) of each significant cluster are indicated by the horizontal extent of grey shaded regions. Results are depicted for **a)** a test for differences in oscillatory power when people were instructed to imagine having audio as opposed to visual experiences, **b)** a test for differences in oscillatory power when people were instructed to meditate as opposed to having imagined audio experiences, and **c)** a test for differences in oscillatory power when people were instructed to meditate as opposed to having imagined audio experiences.

**Figure 6.**
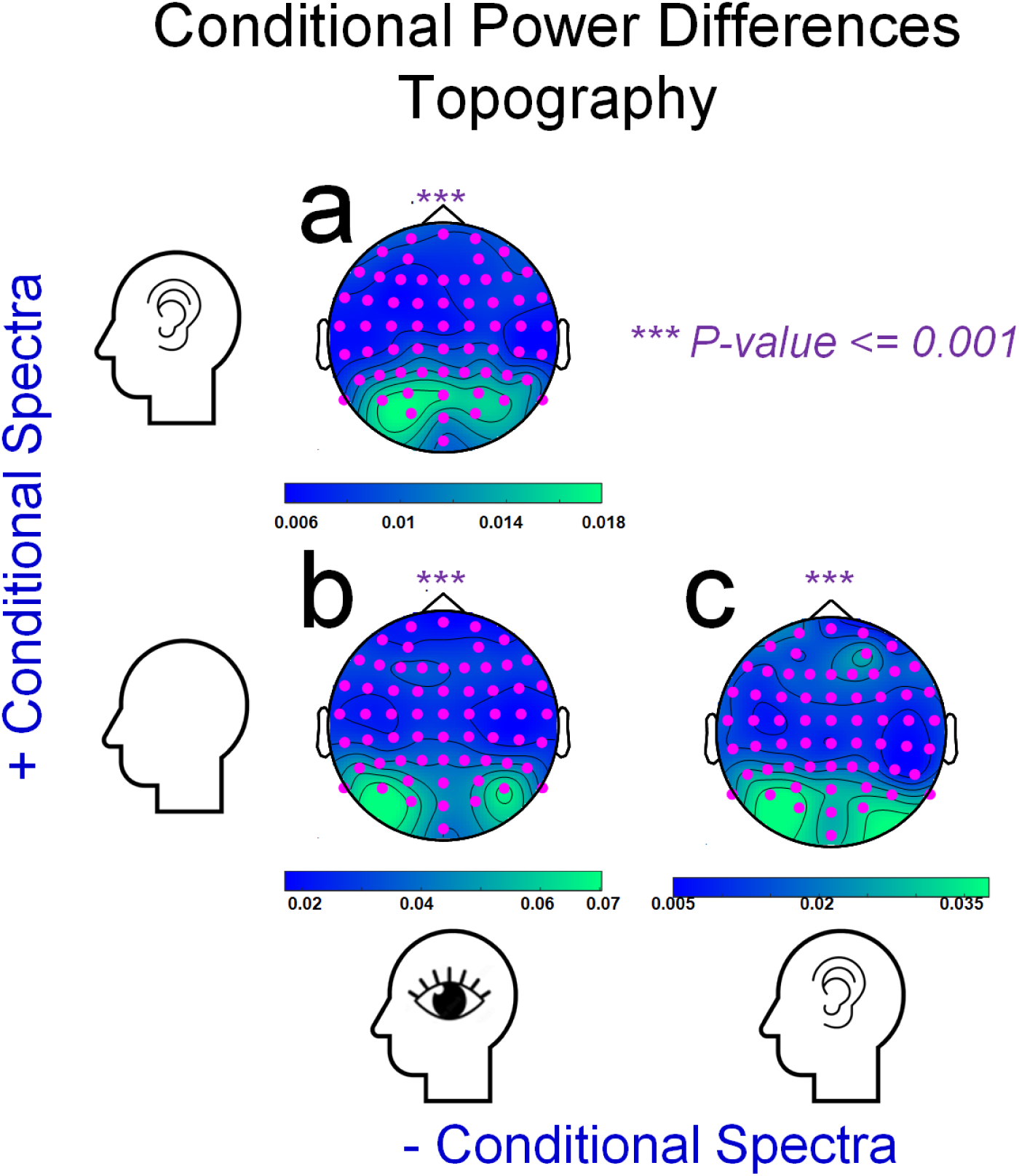
Details are as for Figure 5, with the following exceptions. Panel graphics depict heatmaps of differences in estimates of oscillatory power, recorded by each sensor and averaged across cluster frequencies (as depicted in Figure 5). In each case data recorded by all sensors contributed to these significant cluster tests, as indicated by the distributions of pink dots.

We conducted three cluster tests, to detect differences between spectra calculated from brain activity recorded for each combination of our experimental conditions. These tests were based on paired t-tests, comparing individual conditional estimates of the oscillatory power of brain activity (on average, for each combination of sensor and frequency). Tests with an uncorrected p-value < 0.05 are clustered, based on spatio-frequency proximity, and cluster-level statistics are obtained by summing test statistics and taking the maximum to test significance against a random distribution, obtained via 1000 permutations of the original data [24].

As depicted in Figures 5 and 6, each of these tests was significant (p-values < 0.001), establishing that spectra which describe the brain activity which prevailed during trials for each experimental condition was different (was not interchangeable) with the spectra that prevailed during trials for each of the other two experimental conditions.

For the test for spectra differences, while people attempted to have imagined audio as opposed to visual experiences, differences were detected across a broad range of frequencies (∼0 – 24 Hz) with a prominent alpha band peak (∼9Hz, see Figure 5a). In terms of topography, all sensors contributed to the significant cluster test, but differences were maximal at occipital sensors (see Figure 6a, pink dots mark sensors that contributed to the significant cluster test).

The test for spectra differences, while people attempted to meditate as opposed to having visual experiences, also detected differences across a broad range of frequencies (∼0 – 10 Hz), with a similarly prominent alpha band peak (∼9Hz, see Figure 5b). In terms of topography, again all sensors contributed to the significant cluster, but differences were again maximal at occipital sensors (see Figure 6b).

The test for spectra differences, while people tried to meditate as opposed to having audio experiences, also detected differences across a broad range of frequencies (∼5 – 14 Hz), with a prominent alpha band peak (∼9Hz, see Figure 5c). In terms of topography, differences were again detected across all sensors, but were maximal at occipital sensors (see Figure 6c).

In sum, our cluster tests show that all experimental conditions were associated with distinct spectra, which were reliable across participants. This, however, does not speak to the issue of whether the spectra that describe brain activity as people attempt to imagine having different types of sensory experience can also predict the subjective intensities of different peoples’ imagined experiences. We address this issue via a set of linear support vector regression (SVR) analyses [27].

### Linear Support Vector Regression Analyses

Initial treatment of data for these analyses was as described for cluster tests, including the exclusion of feature outliers. To isolate features of brain activity that were involved in generating imagined sensory experiences, from oscillatory activity that is generic for an individual, we then created conditional difference spectra, by subtracting individual spectra calculated from meditation trials from individual spectra calculated from trials involving imagined audio and imagined visual experiences.

In addition to isolating the features of brain activity that were involved in generating imagined sensory experiences, the calculation of conditional difference scores has the advantage of subtracting out features of brain activity that might be specific to a recording session (i.e. greater overall estimates of oscillatory power, due to less noise from a better EEG cap fit), as opposed to the brain activity that is set in train by the cognitive operations we wish to investigate. To further simplify these analyses, we averaged difference scores across all sensors for each individual. So, the following analyses are a test for predictive relationships between *global* measures of oscillatory power differences (i.e. difference scores averaged across all sensors) and measures of the subjective intensity of imagined experiences.

The first set of SVR analyses involved subjective intensity ratings relating to imagined audio experiences. An audio SVR analysis was conducted for each frequency. The feature space for these was a 44 (participants) X 1 (audio frequency conditional difference scores) space, used to predict subjective audio intensity ratings. SVRs involved 125 iterations of a 4-fold cross validation process. For each iteration, individual participants were randomly assigned to one of four evenly sized (N=11) groups. For each fold, the features and intensity ratings for 3 of the groups (N=33) were used to train a linear regression model, using the Matlab fitsvm command. The resulting model was then used to predict subjective audio intensity ratings of the test group (N=11), based on test group audio EEG features, and a correlation coefficient was calculated between these predicted and the actual test group audio intensity ratings. Resulting R-values were stored for analyses. A matching set of shuffled analyses was also undertaken, wherein all participant intensity ratings had been randomly reassigned prior to the formation of groups – to create a distribution of chance R-values (where there can be no correspondence between EEG features and subjective audio intensity ratings other than chance). Shuffled chance R-values were also stored for comparison with actual R-values.

The 125 iterations of the 4-fold process generate 500 actual and 500 shuffled chance R-values for each frequency (1-40Hz). Bayes Factor t-tests were conducted for each frequency, to determine if there is evidence for a difference between actual and chance R-values, or if there is evidence for these being interchangeable. These are plotted in Figure 7a.

**Figure 7.**
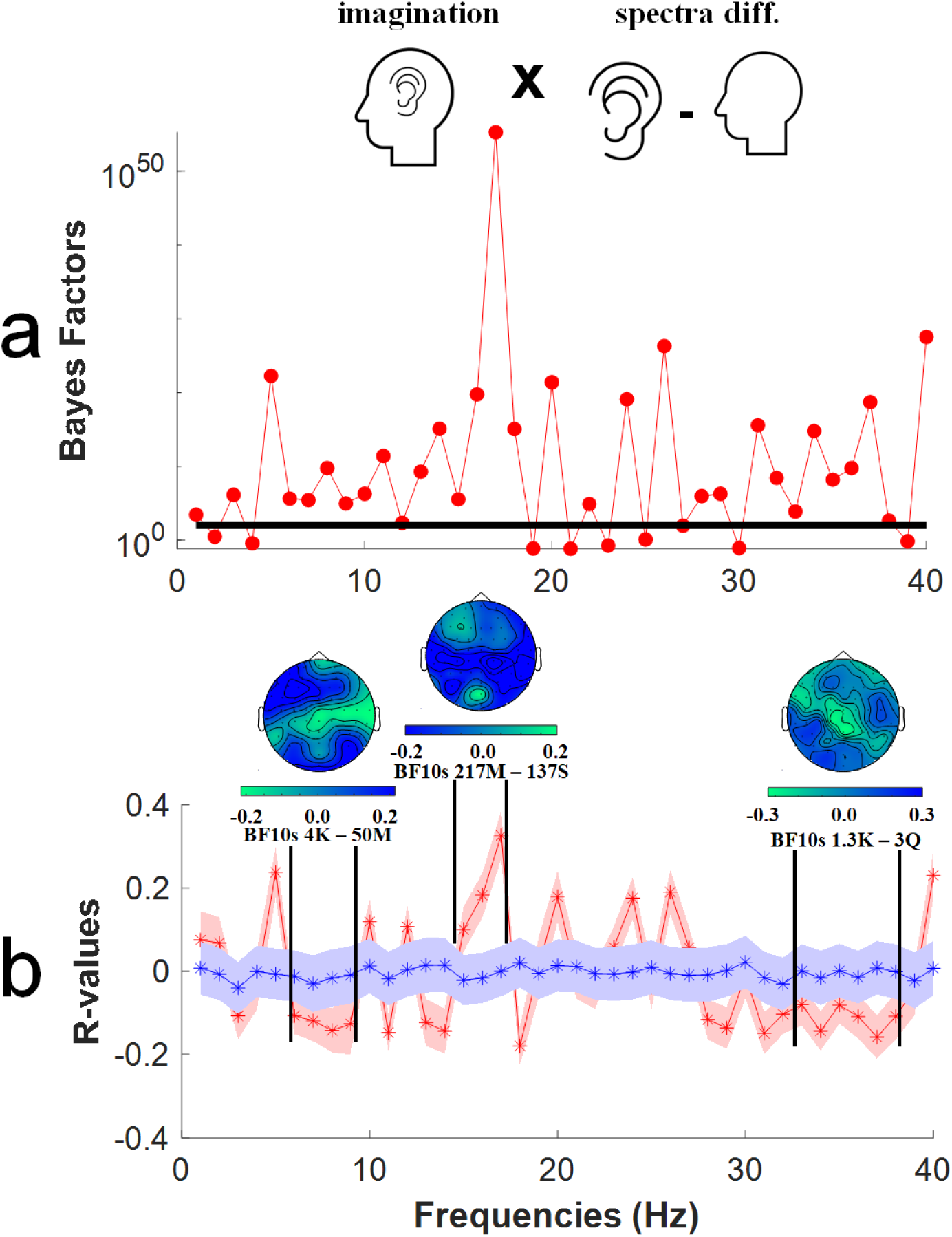
Results of non-parametric support vector regression (SVR) analyses. Analyses were conducted to detect relationships between estimates of oscillatory power averaged across all sensors and individual subjective ratings of imagined audio intensities. **a)** Bayes Factors (Y-axis) as a function of oscillatory frequency (Hz, X-axis). The bold horizontal line marks a BF_10_ = 100, which constitutes extreme evidence for the alternative hypothesis, that there is a predictive relationship between spectra differences and the intensity of imagined audio experiences. **b)** R-values (Y-axis) as a function of oscillatory frequency (Hz, X-axis). The red line marks the average R value of regressions informing the audio SVR analysis, and the blue line marks the average R value of shuffled chance regressions. In each case shaded regions mark +/- 2SEM. Insert heatmaps depict the distribution of R-values across individual sensors, averaged across cluster frequencies. The frequency limits of clusters are depicted by bold vertical black lines. Note that individual sensor difference scores were averaged into a global measure for SVR analyses. See main text for further details.

While the non-parametric treatment of these data provide protection against detecting false positive relationships, further evidence against spurious relationships is provided by clusters of predictive relationships (i.e. in this context, when there are similar predictive relationships across successive oscillation frequencies). In these data, there are three such predictive clusters. Actual (red data) and shuffled chance R-values are plotted in Figure 7b. Here three clusters of predictive frequencies (6 – 9 Hz, 15 – 17 Hz, and 33 – 38 Hz) are highlighted, where evidence for predictive relationships is again overwhelming (6 – 9 Hz minimum BF_10_ > 4,000; 15 – 17 Hz minimum BF_10_ > 217 Million; 33 – 38 Hz minimum BF_10_ > 1,300). Respectively, these clusters of predictive relationships reside within the theta, beta and gamma bands of oscillatory frequencies. Overall, these data show that in this sample of participants estimates of the power of oscillatory activity can be used to predict the subjective intensity of imagined audio experiences.

A second set of visual SVR analyses involved subjective intensity ratings for imagined visual experiences. Details of these analyses are as described for audio SVRs. Visual imagination bayes factors are plotted as a function of test oscillation frequency (1-40Hz) in Figure 8a. To highlight *clusters* of predictive relationships, actual (red data) and shuffled chance R-values are plotted in Figure 8b. Here, three clusters of predictive frequencies (6 – 9 Hz, 15 – 17 Hz, and 33 – 38 Hz) are highlighted, where evidence for predictive relationships is again overwhelming (14 – 16 Hz minimum BF_10_ > 4.7 Billion; 24 – 26 Hz minimum BF_10_ > 3.8 Sextillion; 29 – 32 Hz minimum BF_10_ > 28.5 Million). Respectively, these clusters of predictive relationships reside within the lower, mid and upper beta bands of oscillatory frequencies. These data show that, in our sample of participants, estimates of the global power of oscillatory activity can be used to predict the subjective intensity of imagined visual experiences.

**Figure 8.**
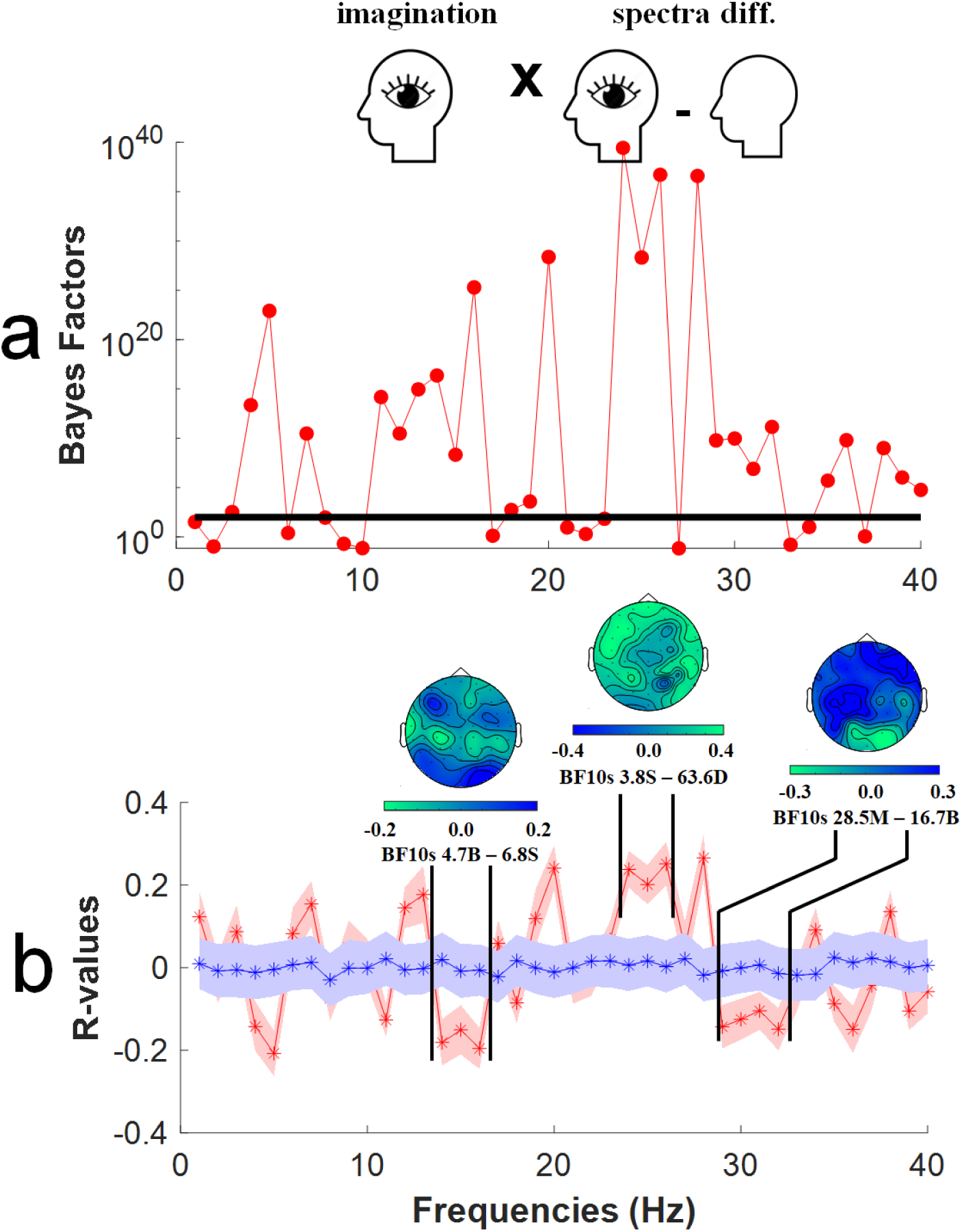
Results of non-parametric support vector regression (SVR) analyses, to detect predictive relationships between estimates of oscillatory power and subjective ratings of imagined visual intensities. All other details are as for Figure 7.

## Discussion

We have found that the subjective intensity of different peoples imagined visual and auditory experiences can be predicted from conditional differences in the power of oscillatory brain activity. For imagined audio experiences, there were clusters of predictive frequencies in the theta, beta and gamma oscillation frequency bands (see Figure 7). For imagined visual experiences, there were clusters of predictive frequencies in the lower, mid and upper beta frequency bands (see Figure 8).

### The validity of subjective ratings of the intensity of imagined sensory experiences

Here, we have primarily been interested in determining if measures of the power of oscillatory brain activity can be used to predict the subjective intensity of different peoples imagined sensory experiences. Our data provide strong support for this notion (see Figures 7 and 8). The relationships we have identified should be examined in further studies, to determine if they generalise to other samples of people. However, the fact that we have been able to detect predictive relationships suggests there is at least some ground truth to the subjective ratings people use to describe the intensity of their imagined experiences, as these ratings could be predicted by the distinct spectra that described different people’s brain activity as they try to imagine having audio and visual experiences.

Subjective ratings are, however, undoubtedly a noisy and imprecise measure of the intensity of imagined sensory experiences. Highlighting this issue, a research student who works in our lab is a profound aphantasic – who not only reports being unable to conjure imagined visual images, but also reports having no inner voice, and an absence of visual or auditory experiences while dreaming. Nonetheless, when this student first completed the VVIQ2 questionnaire [23] her responses were stereotypical, as she was not then aware that other people could have imagined sensory experiences, and so misconstrued the questions as relating to the effort expended and success in remembering facts about sensory experiences. Such issues could perhaps be mitigated by educating participants about the possibility of variable outcomes when different people try to have imagined sensory experiences before they engage in experiments. However, more reliable metrics of the intensity of imagined experiences are desirable in this context (for a related discussion, see [28]).

Attempts to identify tasks where performance is contingent on the subjective intensity of imagined sensory experiences have not always been successful [e.g. 29]. There are, however, some reported exceptions. The probability of detecting visual targets that are subject to binocular suppression can reportedly be enhanced by having people pre-imagine inputs prior to stimulus presentations [9]. Pupil dilations are also reportedly responsive to the brightness of imagined content, in a manner that scales with the subjective intensity of imagined experiences [30]. Finally, people who have vivid imagined visual experiences reportedly experience greater interference when trying to read a briefly presented (and backwardly masked) word when the meaning of that word is incongruent with a background colour [11]. It is possible that one or all of these behavioural measures might provide a more accurate index of the vividness of imagined experiences than subjective questionnaires – but how would we know?

Until now, studies that have strived to identify a behavioural task, where performance is contingent on the intensity of imagined sensory experiences, have only been able to attempt validation by comparing task performance to subjective responses on questionnaires [9,11,29,30]. Our results suggest future studies could attempt validation by comparing task performances with measures of brain activity, and in particular to measures of the power of oscillatory brain activity. The data we have presented might serve as an important guide, as to what spectra of brain activity might be fruitfully targeted by these investigations.

### Could our findings be explained via eye movements?

We don’t think so, but we would not be at all upset if they were.

While all recordings of brain activity were taken while people had closed eyes, it is nonetheless possible that people might have moved their eyes while they engaged in imagining experiences, and that this could have contaminated our measures and predicted the subjective intensity of imagined experiences. It has been thought that people with vivid imaginations are more likely to gaze about their imagined vistas [31], and some people report that they feel this can assist them to have vivid imagined visual experiences [32]. However, evidence for this relationship is tenuous, with multiple report of there being no such relationship [e.g. 33-34] and at least one instance of evidence for an opposite relationship (when imagining static scenes, eye muscle activity was negatively correlated with the subjective intensity of imagined images, see [34]).

In our study we used ICA analyses, implemented in FieldTrip [24], to detect and remove eye and muscle artifacts in pre-processing before data analyses. This approach has been shown to be effective in removing ocular artifacts [35]. We also took the precaution of excluding any sensor trial data that exceeded a range of 250mV from analyses, and neither the spectra or the topography of the predictive relationships we have identified (see Figures 7 and 8) are reminiscent of ocular artifacts [see 35]. In short, we do not believe there is any evidence in our data to suggest that the predictive relationships we have identified can be explained by eye movements.

We would further note, however, that we would welcome such a simple predictive relationship. If people with vivid imaginations were more likely to look about their imagined vistas as they engage in sensory imagination, this could serve as an easily recorded biomarker to predict the subjective intensity of a persons’ imagined experiences. Alas, there is no evidence for that relationship in our data. We would recommend, however, that future studies record electrooculogram (EOG) in addition to EEG – to provide further insight into any interrelationships that might exist between eye movements and the subjective intensity of imagined sensory experiences in different circumstances.

### The intensity of imagined experiences is not predicted by peak alpha band oscillations

Each of our experimental conditions was associated with distinct spectra that described the brain activity recorded during that condition, and these differences were reliable across participants (see Figures 4 and 5). So, the act of imagining auditory experiences, visual experiences, or meditating, each elicited a distinct pattern of brain activity described by different spectra. In each case conditional differences had a prominent peak in the alpha band (∼9Hz, see Figure 4) and was maximal in recordings taken from occipital sensors (see Figure 5). These differences, however, speak merely to the act of having imagined experiences. As can be seen in Figures 7 and 8, this frequency and distribution of oscillatory activity did not predict individual differences in the subjective *intensity* of imagined experiences.

Our findings, regarding peak differences in the alpha band (see Figure 4), are consistent with a number of past investigations that have similarly implicated changes in peak alpha-band oscillations as a marker of when people are engaged in having imagined sensory experiences [e.g. 19-21]. However, when considered in conjunction with our SVR analyses, our findings suggest that these largest conditional differences do not predict the subjective intensity of different people’s imagined experiences. Rather, clusters of frequencies in the theta, beta and gamma bands have predicted the subjective intensities of imagined audio experiences (see Figure 7), and clusters of low, mid and high beta band frequencies predicted the subjective intensities of imagined visual experiences (see Figure 8). Past researchers might have failed to detect these predictive relationships if they were focussed on larger, more obvious conditional differences that are not predictive of the subjective intensity of imagined experiences.

### Topography of predictive oscillations

In terms of the topography of predictive sensors, in our study these encompassed all sensors (as the oscillatory activity informing our SVRs was a global measure). Topographic maps highlighted contributions from frontal, central and posterior sensors (see Figures 7 and 8). This dispersed distribution is broadly consistent with the notion that the oscillatory activity that predicts the subjective intensity of different peoples imagined sensory experiences is driven by coordinated responding from across broad networks of distributed brain regions [36], with key processes originating in frontal brain regions [14, 37]. Our data are also broadly consistent with the premise that the subjective intensity of imagined experiences is governed by the coordinated action of a global neuronal workspace [38].

### Predictive oscillations and established links to cognition

In this study, the core goal was to establish if the power of oscillatory brain activity could predict the subjective intensity of different peoples imagined sensory experiences. Our data suggest this can be done, as there are multiple clusters of predictive oscillatory activity evident in our data. Some of these have established links with cognitive processes that are likely triggered by attempts to have imagined sensory experiences. For instance, in our study, the subjective intensities of both audio and visual imagined experiences were predicted by clusters of beta band frequencies, and these have previously been linked with working memory operations [39-40] and with endogenous top-down perceptual processes [41-42]. Indeed, contemplation of the later findings led to the prediction that mental imagery of sensory events would be associated with beta band oscillations [see 43], and that prediction has found some level of support in our data (see Figures 7 and 8).

There are other predictive relationships in or data, and links between these and cognitive operations are less clear. Indeed, linking these findings to individual cognitive operations might fundamentally misconstrue the true significance of our findings, which may speak to the importance of coordinated responding from across broad networks of brain regions [36], rather than to modular brain regions or cognitive functions. At this point, we would merely note that all the predictive relationships evident in our data should be re-examined in further studies, to establish the generality of our findings.

### Imagined sensory experiences and mental representations

The conceptual background for our study is an historic debate concerning the nature of mental representations. One school of thought has maintained that these are stored exclusively in a language-like propositional (descriptive) format [e.g. 44], whereas another has maintained that sensory representations are characterised by pictorial qualia – by consciously experienced images [e.g. 45]. Our data highlight individual differences, so it seems plausible that for some people (i.e. aphantasics, see [2]) mental representations are purely propositional, whereas the majority of people can conjure imagined sensory experiences that manifest as variably intense, consciously experienced qualia. This conjecture is consistent with the different descriptions aphantasics use to describe how they achieve tasks, as opposed to the majority of people who assert they perform the same tasks by tapping on their ability to conjure mental images [see 46].

### Conclusions

Our results make two important contributions. **1)** They suggest that the spectra of brain activity is a viable metric of the subjective intensity of different peoples imagined sensory experiences. **2)** They suggest that peak alpha band oscillatory brain activity does not predict the subjective intensity of imagined sensory experiences – even though these were the largest, most obvious oscillatory differences triggered by attempts to have imagined sensory experiences. Instead, we identified a number of other clusters of predictive oscillatory frequencies. Our data are consistent with imagined sensory experiences being driven by coordinated responding from across broad networks of distributed brain regions. We believe the predictive oscillatory activity we have identified should be re-examined in future targeted investigations, that also incorporate objective measures of the intensity of imagined audio and visual experiences.

## Acknowledgements

This research was supported by a Discovery Project Grant, funded by the Australian Research Council, awarded to D.H.A. We would like to thank Dr. Amanda Robinson for feedback on a draft version of this manuscript.

We would like to thank Dr. Amanda Robinson for being an all-round awesome person, and for giving us feedback on draft versions of this manuscript.

